# Hierarchical genomic feature annotation with variable-length queries

**DOI:** 10.64898/2026.03.15.711907

**Authors:** Jarno N. Alanko, T. Rhyker Ranallo-Benavidez, Floris P. Barthel, Simon J. Puglisi, Camille Marchet

## Abstract

*K*-mer-based methods are widely used for sequence classification in metagenomics, pangenomics, and RNA-seq analysis, but existing tools face important limitations: they typically require a fixed *k*-mer length chosen at index construction time, handle multi-matching *k*-mers (whose origin in the indexed data is ambiguous) in ad-hoc ways, and some resort to lossy approximations, complicating interpretation. We present HKS, a data structure for exact hierarchical variable-length *k*-mer annotation. Building on the Spectral Burrows– Wheeler Transform (SBWT), a single HKS index is constructed for a specified maximum query length *s*, and supports queries at any length *k* ≤ *s*. HKS associates each *k*-mer with exactly one label from a user-defined category hierarchy, where multi-matching *k*-mers are resolved to their most specific common node in the hierarchy.

We formalize a feature assignment framework that partitions indexed *k*-mers into disjoint sets according to a user-defined category hierarchy. To recover specificity lost to multi-matching and novel *k*-mers, we introduce a hierarchy-aware smoothing algorithm that makes use of flanking sequence context. We validate the approach by assigning each query *k*-mer to a specific chromosome across human genome assemblies, including the T2T-CHM13v2.0 reference as a positive control and two diploid genomes of different ancestries (HG002, NA19185). Smoothing increases overall concordance from ∼81% to ∼97%, with residual errors attributable to known biological phenomena including acrocentric short-arm recombination and subtelomeric duplications. In performance benchmarks against Kraken2, HKS provides comparable query throughput while providing exact, lossless annotation across all *k*-mer lengths simultaneously from a single index. A prototype implementation is available at https://github.com/jnalanko/HKS.

## 1 Introduction

*K*-mer-based methods underpin a wide range of sequence classification tasks in genomics. In pangenomics, colored de Bruijn graphs [19, 25] associate each *k*-mer with the set of genomes or samples in which it appears, enabling fast membership queries across large reference collections [6, 13, 20]. In RNA-seq analysis, pseudoalignment methods such as kallisto [11, 28] assign reads to transcripts based on shared *k*-mer content, with ambiguity resolved through expectation–maximization. In taxonomic classification, tools such as Kraken [36, 35] assign each *k*-mer to the lowest common ancestor (LCA) of the genomes containing it, propagating labels up a taxonomic tree to handle sequences shared across taxa. Although these applications differ in their biological goals, they share a common computational task: assigning *k*-mers to a structured set of labels. Table 1 classifies existing tools along two dimensions: the labeling scheme used, and whether query computation is exact or approximate.

**Table 1.**
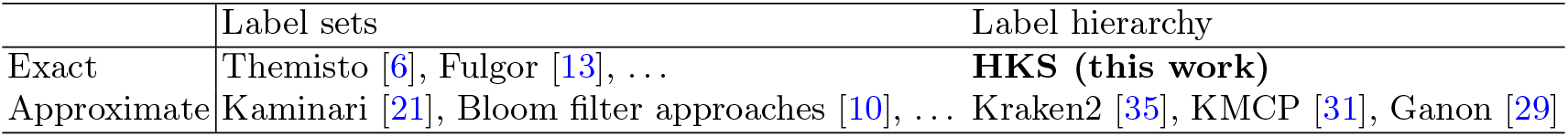
Comparison of *k*-mer labeling paradigms.

Despite this shared foundation, three limitations affect existing *k*-mer-based assignment methods. First, the value of *k* must typically be fixed at index construction time. Short *k*-mers produce more matches but are frequently shared across multiple categories, while long *k*-mers are more specific but fail to match when the query diverges from the index by even a single nucleotide. In practice, this forces users to choose a single compromise value of *k*, or to build and query multiple indexes, requiring results to be coalesced in some way in postprocessing. Second, *k*-mer multi-matching — where a *k*-mer occurs in multiple categories — is handled inconsistently across the literature. Some approaches mask repetitive *k*-mers at query time [8], sacrificing information for simplicity. Others resolve ambiguity through probabilistic models [11, 26] or by propagating categories up a hierarchy [36]. Third, many tools rely on lossy approximations — such as minimizer-based hashing [35] or Bloom filters [10] — that reduce index size and improve speed at the cost of accuracy. No existing tool combines exact hierarchical resolution with the flexibility to query at multiple *k*-mer lengths from a single index.

In this paper, we present HKS, a data structure that addresses these limitations. HKS enables exact *k*-mer-based sequence annotation with respect to a user-defined category hierarchy, which may represent a taxonomy, a set of chromosomes, repeat families, or any other hierarchical organization of genomic labels. We make three contributions:

1. **A feature assignment framework**. We formalize how indexed *k*-mers are partitioned into disjoint sets according to a user-defined category hierarchy. Each *k*-mer that occurs in multiple categories is assigned to the most specific common ancestor of those categories, guaranteeing a unique label per *k*-mer while preserving hierarchical information. This generalizes the LCA strategy used by taxonomic classifiers such as Kraken [36, 35] to any user-defined hierarchy, building on the feature assignment approach introduced in KaryoScope [30].
2. **A variable-length exact index**. Built on the Spectral Burrows–Wheeler Transform (SBWT) [5], HKS assigns each indexed *k*-mer exactly one label from the category hierarchy. The index takes a parameter *s* and supports exact queries for any *k* ≤ *s*, eliminating the need to rebuild for different *k*. Thus, HKS effectively realizes a colored variable-order de Bruijn graph [9], a data structure that has long languished as a theoretical curiosity.
3. **A hierarchy-aware smoothing algorithm**. To recover specificity lost to multi-matching and novel *k*-mers, we introduce a post-processing step, extending the approach of KaryoScope [30], that uses flanking sequence context and the category hierarchy to reassign non-specific *k*-mers to more informative labels.

Together, these components produce a positional annotation of query sequences: each *k*-mer in the query receives a label, and consecutive *k*-mers with the same label are merged into intervals, yielding a segmentation of the query by feature. This contrasts with tools that assign a single label to an entire query sequence, and enables the detection of boundaries between features within a single sequence. For example, when chromosomes are used as categories, a query sequence originating from a single chromosome is annotated almost entirely with one chromosome label, while a translocation manifests as adjacent segments assigned to different chromosomes. We illustrate this with a positional annotation of the RPE1 cell line genome assembly [34], which reveals a known translocation between chrX and chr10 (Supplementary Figure S1). Beyond large-scale rearrangements, the positional resolution of HKS annotations can reveal subtler patterns of inter-chromosomal sequence sharing, such as recombination between acrocentric short arms or subtelomeric segmental duplications, that would be obscured by whole-sequence classification.

We validate HKS by querying three human genome assemblies against a T2T-CHM13v2.0 [27] chromosome index and assigning each *k*-mer to a specific chromosome. Using each assembly’s chromosome of origin as ground truth, we show that pre-smoothed annotations achieve near-perfect accuracy (99.8%) but leave ∼19% of *k*-mers unresolved to a specific chromosome. After smoothing, chromosome-specific concordance increases from ∼81% to ∼97%, with residual errors attributable to well-characterized biological phenomena rather than algorithmic failures.

We benchmark the computational performance of HKS against Kraken2, a widely-used tool for hierarchical *k*-mer classification. Unlike Kraken2, which requires a separate index for each value of *k* and relies on lossy minimizer-based approximations, HKS supports exact queries at any *k* ≤ *s* with *s* fixed at the time of indexing. Despite this generality, HKS provides comparable query throughput to Kraken2 across all tested *k*-mer lengths, and when Kraken2 is configured for exact matching (*m* = *k*), HKS is both faster and produces a smaller index. The HKS implementation is available at https://github.com/jnalanko/HKS.

## 2 Preliminaries

In this section we define notation, terminology, and basic concepts used throughout. We use the usual array notation for strings. We consider a string *S* of length |*S*| = *n* as a sequence of *n* symbols *S*[0..*n* − 1] = *S*[0]*S*[1] … *S*[*n* − 1] drawn from an alphabet Σ = {A, C, G, T} and *σ* = |Σ| = 4. The empty string is denoted *ϵ* and |*ϵ*| = 0. We use 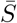 to denote the reverse complement of *S*. We denote with *S*^*k*^ the string *S* repeated *k* times. The *substring* of *S* starting at symbol *i* and ending at symbol *j* is denoted *S*[*i*..*j*]. We also use the half-open interval notations *S*(*i*..*j*] = *S*[*i* + 1..*j*] and *S*[*i*..*j*) = *S*[*i*..*j* − 1]. A *prefix* is a substring starting at position 0 and a *suffix* is a substring ending at position *n* − 1. The *colexicographic order* of two strings corresponds to the lexicographic order of their reverse strings. A *k*-mer is a (sub)string of length *k*.

### 2.1 Spectral Burrows-Wheeler transform

The set of distinct *k*-mers occurring in a string *T* is referred to as the *k-spectrum* of *T*.

#### Definition 1

*(k-Spectrum)* The *k*-spectrum of a string *T*, denoted with *S*_*k*_(*T*), is the set of all distinct *k*-mers in the string *T* ^*′*^ = $^*k*^*T* . The *k*-spectrum *S*_*k*_(*T*_1_, …, *T*_*m*_) of a set of *m* strings *T*_1_, …, *T*_*m*_ is the union 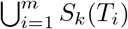

The dollar prefixes are for technical convenience to ensure there is a *k*-mer ending at every position in *T* . By way of illustration, consider the string *T* = TAGCAAGCACAGCATACAGA, for which we have *S*_3_(*T*) = *{*AAG, ACA, AGA, AGC, ATA, CAA, CAC, CAG, CAT, GCA, TAC, TAG $$$, $$T, $TA*}*.

In this paper the primary operation we aim to support is the *k-mer lookup* query, defined as follows:

#### Definition 2

*(k-mer lookup query)* Given an input string *P* of length *k*, a *k*-mer lookup query returns the colexicographic rank of *P* in *S*_*k*_(*T*), or ⊥ if *P*∉ *S*_*k*_(*T*).

For example, the *k*-mers from the 3-spectrum *S*_3_(*T*) above, listed in colexicographical order are as follows: $$$, CAA, ACA, GCA, AGA, $TA, ATA, CAC, TAC, AGC, AAG, CAG, TAG, $$T, CAT. Consequently, the lookup query returns the zero-based rank 4 for AGA, and ⊥ for TTT.

The Spectral Burrows-Wheeler transform (SBWT) [5] transforms a *k*-spectrum into a sequence of subsets of the alphabet. Conceptually simple operations on the SBWT subset sequence enable efficient *k*-mer lookup queries.

#### Definition 3

*(Spectral Burrows-Wheeler Transform)* Let *S*_*k*_ be a *k*-spectrum and let 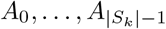 be the colexicographically sorted elements of *S*_*k*_. The SBWT is the sequence of sets of characters 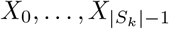 with *X*_*i*_ ⊆ Σ such that *X*_*i*_ = ∅ if *i >* 0 and *A*_*i*_[1..*k*) = *A*_*i*−1_[1..*k*), otherwise *X*_*i*_ = *{c* ∈ Σ | *A*_*i*_[1..*k*)*c* ∈ *S*_*k*_*}*.

The SBWT of the running example is the following sequence of sets: *{*T*}, {*G*}, {*A,C,G,T*}*, ∅, ∅, *{*C,G*}*, ∅, *{*A*}*, ∅, *{*A*}, {*A,C*}*, ∅, ∅, *{*A*}, {*A*}*.

The sets in the SBWT represent the labels of outgoing edges in the node-centric de Bruijn graph, such that only outgoing edges from *k*-mers that have a different suffix of length *k* − 1 than the preceding *k*-mer in the colexicographically sorted list are included. This gives the SBWT sequence an important property, namely that ∑|*X*_*i*_| = *S*_*k*_(*T*) − 1. That is, the sum of the set sizes in the SBWT is exactly one less than the number of *k*-mers in the spectrum, allowing the entire spectrum to be efficiently encoded by the transform. Indeed, one can think of the SBWT as encoding the set of *k*-mers *S*_*k*_(*T*) as a colexicographically sorted list. In the context of *k*-mer lookup, the SBWT allows us to navigate that ordered list via the operation *ExtendRight*, defined as follows.

#### Definition 4

*(ExtendRight)* Let [*s, e*]_*α*_ be the *colexicographic interval* of string *α* with |*α*| *< k*, where *s* and *e* are respectively the colexicographic ranks of the smallest and largest *k*-mer in *S*_*k*_ that have the substring *α* as a suffix. *ExtendRight*([*s, e*]_*α*_, *c*) denotes the *right extension* of the interval [*s, e*]_*α*_ with a character *c* ∈ Σ. This outputs the interval [*s*^*′*^, *e*^*′*^]_*αc*_, or ⊥ if no such interval exists.

As explained in [5], *ExtendRight* can be answered via two *subset rank* operations on the SBWT. We avoid a deeper treatment of this operation here, because it is not essential for understanding the methods of this paper, and refer the reader to the growing literature on the SBWT [5, 2, 3, 4] for further details.

### 2.2 Longest common suffix array and matching statistics

We augment the SBWT with a data structure called the longest common suffix (LCS) array [3] that stores the lengths of the longest common suffixes of adjacent *k*-mers in colexicographical order (we give a precise definition below). The LCS array allows us to support so-called *left contraction* queries: given an interval [*s, e*] in the colexicographical ordering of the *k*-mers of the spectrum containing all the *k*-mers that share a suffix *X* of length *k*^*′*^ ∈ (0, *k*] and a contraction point *t < k*^*′*^, *ContractLeft*([*s, e*], *t*) operation returns the interval [*i*^*′*^, *j*^*′*^] containing all the *k*-mers having *X*[*t*..*k*^*′*^] as a suffix. Left contractions enable streaming *k*-mer search with the SBWT [4, 2].

#### Definition 5

*(Longest common suffix* (LCS) *array) Let {T*_0_, …, *T*_*m*−1_*} be a set of strings and let x*_*i*_ *denote the colexicographically i-th k-mer of S*_*k*_(*T*_0_, …, *T*_*m*−1_). *The LCS array is an array of length* |*S*_*k*_(*T*_0_, …, *T*_*m*−1_)| *s*.*t. LCS*[0] = 0 *and for i >* 0, *the value of LCS*[*i*] *is the length of the longest common suffix of k-mers x*_*i*_ *and x*_*i*−1_.

The SBWT and LCS array can be used to compute the *k-bounded matching statistics* of a query string *Q* with respect to the *k*-mer spectrum indexed by the SBWT.

#### Definition 6

*(k-bounded matching statistics). The k-bounded matching statistics M*_*Q*_ *of a query Q against a k-mer spectrum is the sequence of* |*Q*| *pairs* (*p*_*i*_, *ℓ*_*i*_) *such that ℓ*_*i*_ *is the length of the longest suffix of Q*[0..*i*] *that is a suffix of at least one k-mer in the indexed spectrum. If ℓ*_*i*_ = 0, *then we define p*_*i*_ = −1. *Otherwise, p*_*i*_ *is the rank of the colexicographically smallest k-mer having Q*(*i* − *ℓ*_*i*_, *i*] *as a suffix*.

Alanko et al. [4] show how *ExtendRight* and *ContractLeft* operations on the SBWT and LCS, allow *M*_*Q*_ to be computed efficiently in *Θ*(|*Q*|) operations in a streaming fashion.

### 2.3 Feature assignment

In colored de Bruijn graph approaches such as Themisto [6] and others [13, 20], each *k*-mer is associated with a color set encoding its presence or absence across a collection of samples. In HKS, by contrast, each indexed *k*-mer is associated with exactly one label. This approach builds on KaryoScope [30], which introduced the concept of partitioning *k*-mers into disjoint features using a category hierarchy for chromosome-scale annotation of genome assemblies. To achieve this, we introduce a framework based on four concepts — *categories, category hierarchies, features*, and *feature sets* — that resolves *k*-mers shared across multiple categories into a disjoint labeling.

*Category*. A *category* is a label assigned to indexed *k*-mers according to some criterion — such as `chr1` and `chr2` when labeling by chromosome identity, or `SINE` and `LINE` when labeling by repeat classification. A given *k*-mer may belong to multiple categories; for instance, a *k*-mer that occurs on both `chr13` and `chr21`.

*Category hierarchy*. Categories are arranged in a user-defined tree: leaves are categories, and internal nodes group related ones (the categories in their subtree). For example, in chromosome-based labeling, the acrocentric chromosomes (`chr13`, `chr14`, `chr15`, `chr21`, `chr22`) share a common internal node (Figure 1).

**Fig. 1.**
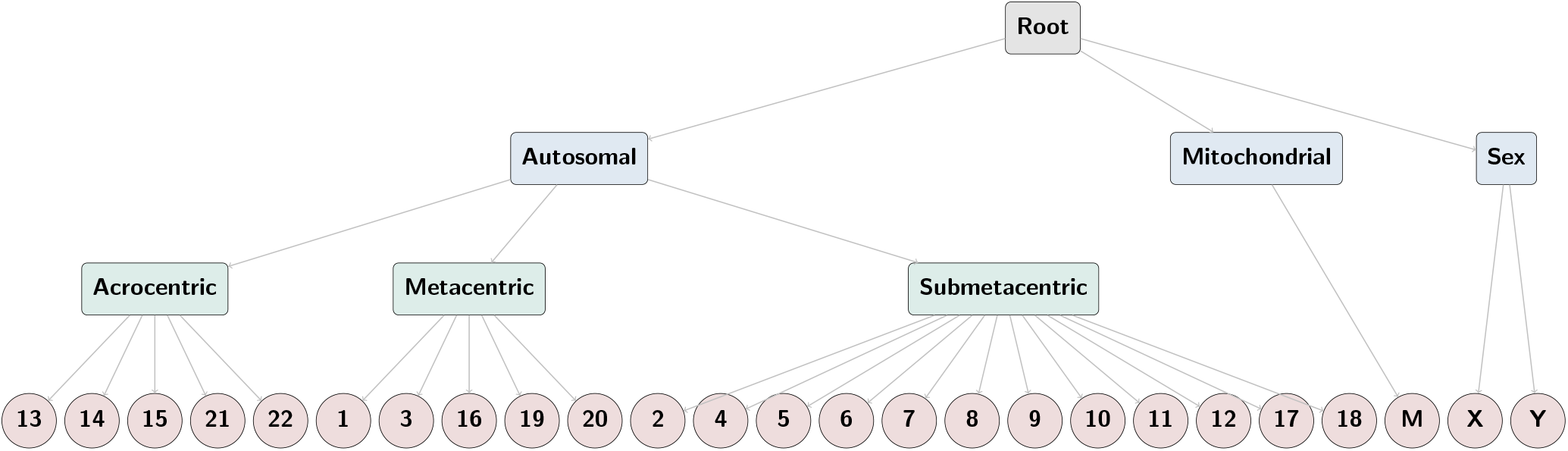
The chromosome category hierarchy used in this work. Leaves represent individual chromosomes (categories) and internal nodes represent groups of related categories. A *k*-mer shared between two or more categories is assigned to the most specific node that encompasses all categories in which it occurs, producing a multigroup feature at that node.

*Feature*. A *feature* is a disjoint set of *k*-mers derived from the category hierarchy. While categories may overlap, features do not: each indexed *k*-mer belongs to exactly one feature. *Category-specific* features contain *k*-mers unique to a category, while *k*-mers shared by multiple categories are assigned to a *multigroup* feature at the most specific node covering them (e.g., a *k*-mer shared by `chr13` and `chr21` belongs to the acrocentric multigroup feature).

*Feature set*. A *feature set* is the complete labeling system comprising a set of categories, a category hierarchy, and the resulting features. Each feature set defines a labeling of the indexed *k*-mers according to a given criterion, such that each *k*-mer is assigned to exactly one feature. A HKS index may incorporate multiple feature sets simultaneously; in this case, each queried *k*-mer receives one feature label per feature set. In this work, we use two feature sets: one based on chromosome identity and one based on repeat classification.

#### Assignment of *k*-mers to features

For a given feature set, feature assignment proceeds in two stages. First, each position in the indexed sequences is assigned to exactly one category. Second, each distinct *k*-mer is assigned to a feature using the category hierarchy as described above. In the first stage, each *k*-mer is assigned to the category of its starting position; thus, if a *k*-mer spans the boundary between two categories, it is assigned to the category in which it starts. Because the same *k*-mer may occur at multiple positions, a distinct *k*-mer may be associated with multiple categories, which are then resolved to a single feature via the category hierarchy. Any query *k*-mer absent from the index is left unassigned.

#### Feature sets for the T2T-CHM13v2.0 human reference genome

In this work, we apply the above framework using two feature sets derived from the T2T-CHM13v2.0 human reference genome: a repeat feature set and a chromosome feature set.

*Repeat feature set*. Repeat categories were derived from the RepeatMasker [32] v4.1.2p1 annotation of T2T-CHM13v2.0, generated with the RMBlast search engine and the Dfam 3.3 repeat library augmented with T2T-derived repeat models [17]. RepeatMasker assigns each annotated interval a repeat family classification. We consolidate these annotations into 13 top-level repeat categories: `LINE`, `SINE`, `LTR`, `DNA`, `RC`, `Retrotransposon`, `rRNA`, `scRNA`, `snRNA`, `tRNA`, `D20S16`, and `Unknown`, together with a `nonrepeat` category for unannotated positions. Because RepeatMasker annotations may overlap, we resolve overlaps using a predefined priority order (Supplementary Table S2) so that each genomic position receives exactly one repeat category. The repeat category hierarchy follows the Dfam transposable element classification [33] (Figure 2).

**Fig. 2.**
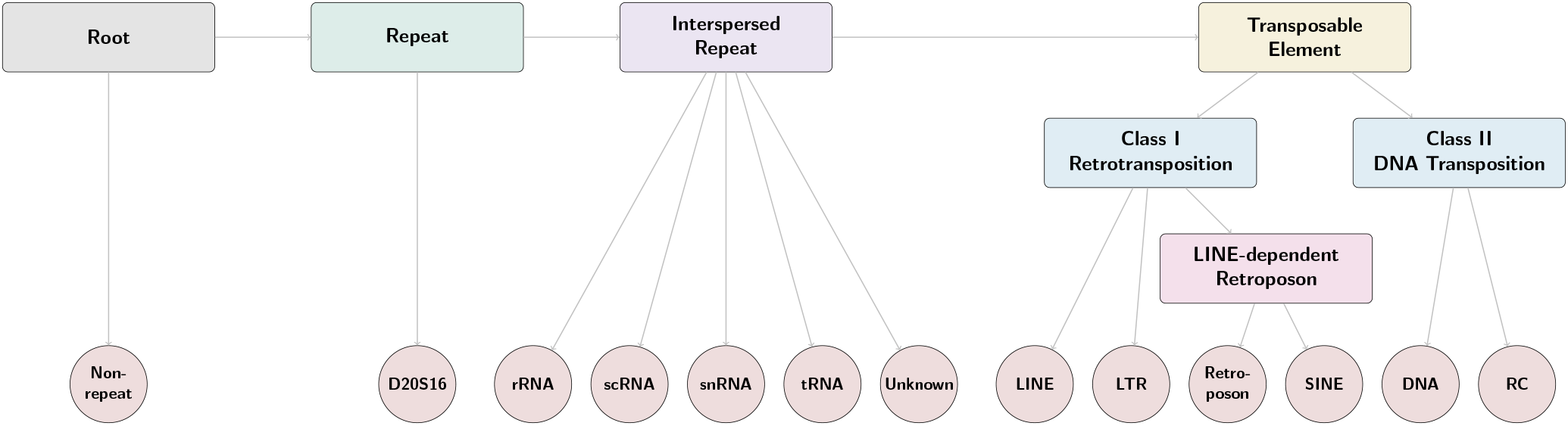
The repeat hierarchy used in this work.

*Chromosome feature set*. Chromosome categories correspond to the T2T-CHM13v2.0 autosomes (`chr1` –`chr22`), sex chromosomes (`chrX`, `chrY`), and mitochondrial genome (`chrM`). These categories are inherently non-overlapping, since each genomic position belongs to exactly one chromosome. The chromosome category hierarchy groups chromosomes by morphology — for example, the acrocentric chromosomes form a shared internal node (Figure 1).

## 3 HKS

Our index, HKS, is constructed from a collection 𝒞 = {*T*_1_, *T*_2_, … *T*_*t*_} of *t* sequences and a hierarchy ℋ, a labeled tree of *m* nodes expressing relationships between the sequences. The sequences are represented as leaves in the hierarchy. Figures 1 and 2 provide two examples of such hierarchies. We emphasise, however, that HKS is more general and can be constructed on any set of genomic sequences with an associated hierarchy, such as microbial strains organized by a phylogenetic tree.

### 3.1 Index Construction

The HKS index consists of four main structures. The first of these is the SBWT for 𝒞. In particular, the SBWT is constructed from the *s*-spectrum 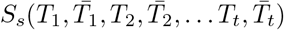, to ensure that both orientations of every *s*-mer in are 𝒞 present in the index. Further, we assume that all *s*-mers belong to Σ^*s*^, that is, they contain only characters from the DNA alphabet—others are discarded. We also store the longest common suffix (LCS) array, denoted *L*, associated with the SBWT, which enables efficient range queries over colexicographically adjacent *s*-mers.

We also store the hierarchy ℋ, which we preprocess for fast lowest-common-ancestor (LCA) queries: the LCA of two nodes *u* and *v*, denoted *LCA*(*u, v*) is the deepest node (furthest from the root) that is an ancestor of both *u* and *v* (with *LCA*(*u, u*) being *u*). LCA is widely studied and has many efficient solutions (see, e.g., [7]). Because in our application ℋ is fairly small (up to 32 nodes, see Fig. 1), we simply precompute and store a table containing the LCA of all *m*^2^ possible pairs of nodes, which allows LCAs to be answered in *O*(1) time. For larger hierarchies, adding a more general solution would be straightforward, but is left as future work.

The final component, denoted *V*_*s*_, is a mapping from each *s*-mer of the SBWT to a node in the hierarchy, ℋ. In particular, let *x*_*i*_ be the *s*-mer (in the SBWT) that has rank *i* in colex order. Then *V*_*s*_[*i*] = *v*_*i*_ is the lowest (furthest from the root) node in ℋ under which all sequences containing *x*_*i*_ occur. That is, *V*_*s*_[*i*] is the LCA in ℋ of all sequences containing *x*_*i*_.

The main task in the construction of the HKS index is to construct *V*_*s*_ from *C* and its SBWT and LCS array. This is accomplished by processing each sequence *T*_*i*_ ∈ *C* in turn and performing an *s*-mer lookup for every *s*-mer contained in *T*_*i*_ using the SBWT. As this series of lookups proceeds, *V*_*s*_ is updated to be the LCA in ℋ of all sequences containing the *s*-mer processed so far. Note that these *s*-mer lookups are always successful, because we are using the SBWT of *C*, of which *T*_*i*_ is a part. Initially, every element of *V*_*s*_[*i*] is set to a special symbol ⊥, to indicate that no mapping for the *s*-mer yet exists. At a generic step in construction, we look up *s*-mer *T*_*i*_[*j*..*j* + *s*− 1] using the SBWT and determine its colex rank, say *r*. If *V*_*s*_[*r*] = ⊥ we set *V*_*s*_[*r*] = *i* (i.e. the leaf of ℋ that corresponds to sequence *T*_*i*_) and continue to lookup the next *s*-mer *T*_*i*_[*j* + 1..*j* + *s*]. Otherwise, we update *V*_*s*_[*r*] to be the LCA of nodes *V*_*s*_[*r*] and *i*. The lookup process is implemented by streaming the *s*-bounded matching statistics (Def. 6) to obtain in succession the colex rank of every *s*-mer of *T*_*i*_. Only the SBWT, LCS, LCA structure for ℋ, and *V*_*s*_ are held in memory, with the sequences *T*_*i*_ being streamed from disk.

### 3.2 Query Algorithm

Querying takes as input a query sequence *Q* of length *q* = |*Q*| and a *k*-mer length, with *k* ≤ *s*, where *s* was the *s*-mer length that was used for indexing. The output is a sequence *B* of *q* node labels, such that *B*[*i*] is the lowest node in that contains *k*-mer *Q*[*i*..*i* + *k* − 1], or, if the *k*-mer does not exist in the indexed collection, then *B*[*i*] = ⊥, a special “no node” label. At a high level, labeling of the query is achieved by performing *k*-mer lookups at every position in *Q* and then mapping these into appropriate nodes in the hierarchy. In our application, *Q* is a chromosome, and so a *query run* typically involves millions of *k*-mer lookups.

In resolving a query run, the SBWT is loaded into memory, together with the *L*, and *V*_*s*_ arrays. We then *prime* the index for the desired length *k*-mer. Priming involves transforming *V*_*s*_ into *V*_*k*_ by scanning the LCS array *L* to find *k-mer groups*, maximal segments of the colexicographically ordered *s*-mer spectrum that share the same *k*-length suffix. More precisely, the colexicographical interval [*i, j*) is a *k*-mer group iff *L*[*i*] *< k, L*[*j*] *< k*, and *L*[*i*^*′*^] ≥ *k*, for all *i*^*′*^ ∈ [*i* + 1..*j*). For each *k*-mer group [*i, j*) we compute the LCA in the hierarchy of all adjacent pairs of *s*-mers it contains. Let this LCA be *v*. We then set *V*_*k*_[*i*^*′*^] = *v* for all *i*^*′*^ ∈ [*i*..*j*). *V*_*k*_ is constructed this way in *O*(*n*) time by scanning and overwriting *V*_*s*_. Just as *V*_*s*_ provides a mapping from the colex rank of an *s*-mer to the node in ℋ under which all sequences containing that appear, so *V*_*k*_ provides a mapping from a *k*-mer to the LCA in of all ℋ leaves containing that *k*-mer.

We then use the SBWT and LCS array to compute the *s*-bounded matching statistics (Def. 6) *M*_*Q*_. Note that the same primed index can be used for multiple *Q*. Having computed *M*_*Q*_ = (*p*_0_, *ℓ*_0_), …, (*p*_|*Q*|−1_, *ℓ*_|*Q*|−1_), we then use *V*_*k*_ to turn *M*_*Q*_ into a sequence of *k*-mer nodes from the hierarchy. In particular, label *B*[*i*] is determined as follows. Let *j* = *i* + *k* − 1 be the ending position of the *i*-th *k*-mer. If *ℓ*_*j*_ *< k* then *B*[*i*] = ⊥ because *ℓ*_*j*_ *< k* implies *k*-mer *Q*[*i*..*i* + *k*) does not appear in the indexed collection. Otherwise (*ℓ*_*j*_ ≥ *k*) we set *B*[*i*] = *V*_*k*_[*p*_*j*_]. This is correct because *ℓ*_*j*_ ≥ *k* implies *k*-mer *Q*[*i*..*i* + *k*) is the suffix of the *s*-mer with colexicographic rank *p*_*j*_, and we constructed *V*_*k*_ to point to the LCA of sequences containing that *k*-mer.

Two comments are in order. First, note that if *k* = *s* then there is no need to prime the index—we can just operate on *V*_*s*_ directly. Further, note that the priming step that constructs *V*_*k*_ from *V*_*s*_ could be avoided and each *k*-mer LCA resolved on demand, for example by scanning ranges of *V*_*s*_ for each element of *M*_*Q*_ and issuing LCA queries on ℋ, or by appropriately preprocessing *V*_*s*_ and storing a data structure along with the other components of the HKS index. However, over a long sequence of queries, such as our query runs contain, the above conversion of *V*_*s*_ into *V*_*k*_ pays off, being fast to perform, and subsequently allowing faster LCA query resolution for the *k* specified.

### 3.3 Post-processing by smoothing

In practice, contiguous stretches of category-specific *k*-mers are often interrupted by short runs of multi-matching or unassigned (novel) *k*-mers, for example due to SNPs or small indels between the query and the indexed sequences. To recover precise feature assignments in these regions, we apply a sequence-context-aware smoothing procedure, extending the approach of KaryoScope [30], that uses the assignments of neighboring *k*-mers. The algorithm identifies windows exhibiting a *specific* → *general* → *specific* pattern in the category hierarchy, corresponding to regions where specificity temporarily decreases and then recovers.

Window detection proceeds via a two-phase scan. Starting from a left anchor interval, the *ancestor phase* scans rightward, accepting intervals whose features are ancestors of the left anchor’s feature in the hierarchy while skipping unrelated intervals on other branches. The phase terminates when specificity increases, when the distance between accepted intervals exceeds a threshold (`max_gap`), or when a skipped unrelated interval would create an overlapping window. The *descendant phase* then continues rightward from the last accepted interval (the turning point), accepting intervals whose features are descendants of the turning point’s feature, under symmetric termination conditions. The final accepted interval becomes the right anchor. See pseudocode in Supplementary Algorithm S1.

For each detected window, the algorithm computes the lowest common ancestor in the hierarchy of the left and right anchor features and reassigns all interior accepted intervals to this node. This replaces less informative multi-matching or novel assignments with the more specific feature supported by the flanking context. The procedure iterates until convergence, typically within one or two passes.

Novel *k*-mers smoothed to a feature more specific than the root are reported under that feature; those smoothed only to the root remain unassigned (novel). The `max_gap` parameter (set to 1000 bp) was chosen by evaluating concordance between smoothed annotations of the HG002 assembly [16] and corresponding reference annotations.

## 4 Results

The data structure and query algorithms were implemented in the Rust programming language. The implementation is available at https://github.com/jnalanko/HKS. The central SBWT data structure is from the `sbwt` crate [1].

### 4.1 Validation with chromosome-specific assignments

We benchmarked the HKS chromosome feature set by evaluating the concordance of chromosome-specific *k*-mer assignments against truth labels derived from chromosome-scale genome assemblies. We compared pre-smoothed (directly from the index) and smoothed (context-aware) labels across three samples: T2T-CHM13v2.0 (haploid positive control), HG002 (diploid Ashkenazi male), and NA19185 (diploid Yoruba female). NA19185 was selected as a genetically divergent sample from T2T-CHM13v2.0 with high assembly quality (35/46 T2T scaffolds) from HPRC Release 2 [18, 22], to maximize sensitivity to reference bias effects.

For each assembly, the truth chromosome label for a given *k*-mer was determined by the chromosome on which it resides. Seven acrocentric p-arm contigs in the NA19185 assembly could not be confidently assigned to a chromosome of origin (Supplementary Table S1). We excluded them from the benchmark (32,393,158 *k*-mers, 0.53% of total).

We computed three primary metrics. Classification rate is the fraction of *k*-mers assigned to any chromosome-specific label, as opposed to a non-specific label, measuring HKS’s ability to resolve *k*-mers to a specific chromosome. Accuracy is the fraction of chromosome-specific *k*-mers assigned to the correct chromosome. Overall concordance is the fraction of all *k*-mers that are correctly assigned, equal to the product of classification rate and accuracy. We also computed adjusted versions of accuracy and concordance that exclude biologically expected mismatches (inter-acrocentric chromosome swaps and sex chromosome homology).

Across all samples, smoothing increases overall concordance by 16-17 percentage points: CHM13 83.5% → 100.0%, HG002 81.0% → 96.8%, NA19185 80.8% → 96.9% (Table 2, Supplementary Figure S2). This improvement is primarily driven by the classification rate increasing from ∼ 81%-84% to ∼ 97%-100%, as the smoothing algorithm resolves non-specific *k*-mers into chromosome-specific assignments using context from neighboring *k*-mers.

**Table 2.** Chromosome feature set concordance across three genome assemblies. Classification rate, accuracy, and concordance are shown for pre-smoothed (directly from the HKS index) and smoothed (context-aware) annotations. Adjusted metrics exclude biologically expected mismatches from inter-acrocentric chromosome swaps and sex chromosome homology.

| Sample | Condition | Raw |  |  | Adjusted |  |
| --- | --- | --- | --- | --- | --- | --- |
|  |  | Class. rate | Accuracy | Concordance | Accuracy | Concordance |
| CHM13<br>(reference) | Pre-smoothed | 83.5% | 100.0% | 83.5% | 100.0% | 83.5% |
|  | Smoothed | 100.0% | 100.0% | 100.0% | 100.0% | 100.0% |
| HG002<br>(Ashkenazi) | Pre-smoothed | 81.1% | 99.8% | 81.0% | 99.9% | 81.0% |
|  | Smoothed | 97.6% | 99.2% | 96.8% | 99.9% | 97.5% |
| NA19185<br>(Yoruba) | Pre-smoothed | 81.0% | 99.8% | 80.8% | 99.9% | 80.9% |
|  | Smoothed | 97.4% | 99.5% | 96.9% | 99.8% | 97.2% |

Presmoothed accuracy is near perfect—99.8% for both HG002 and NA19185, and 100% for CHM13. Smoothing also compensates for reference bias: NA19185 has 3.4% novel *k*-mers (absent from the T2T-CHM13v2.0 index) compared to 3.0% for HG002 and 0.0% for CHM13, reflecting ancestry-specific sequence (Figure 3). After smoothing, these novel *k*-mers are resolved to chromosome-specific assignments using flanking context, reducing the novel fraction to 0.0% for all samples. Post-smoothing accuracy remains above 99% (99.2% for HG002, 99.5% for NA19185). The slight decrease reflects the trade-off of resolving ambiguous *k*-mers; however, this trade-off is highly favorable: for NA19185, smoothing correctly resolves ∼971M additional *k*-mers while introducing only ∼17.6M new errors.

**Fig. 3.**
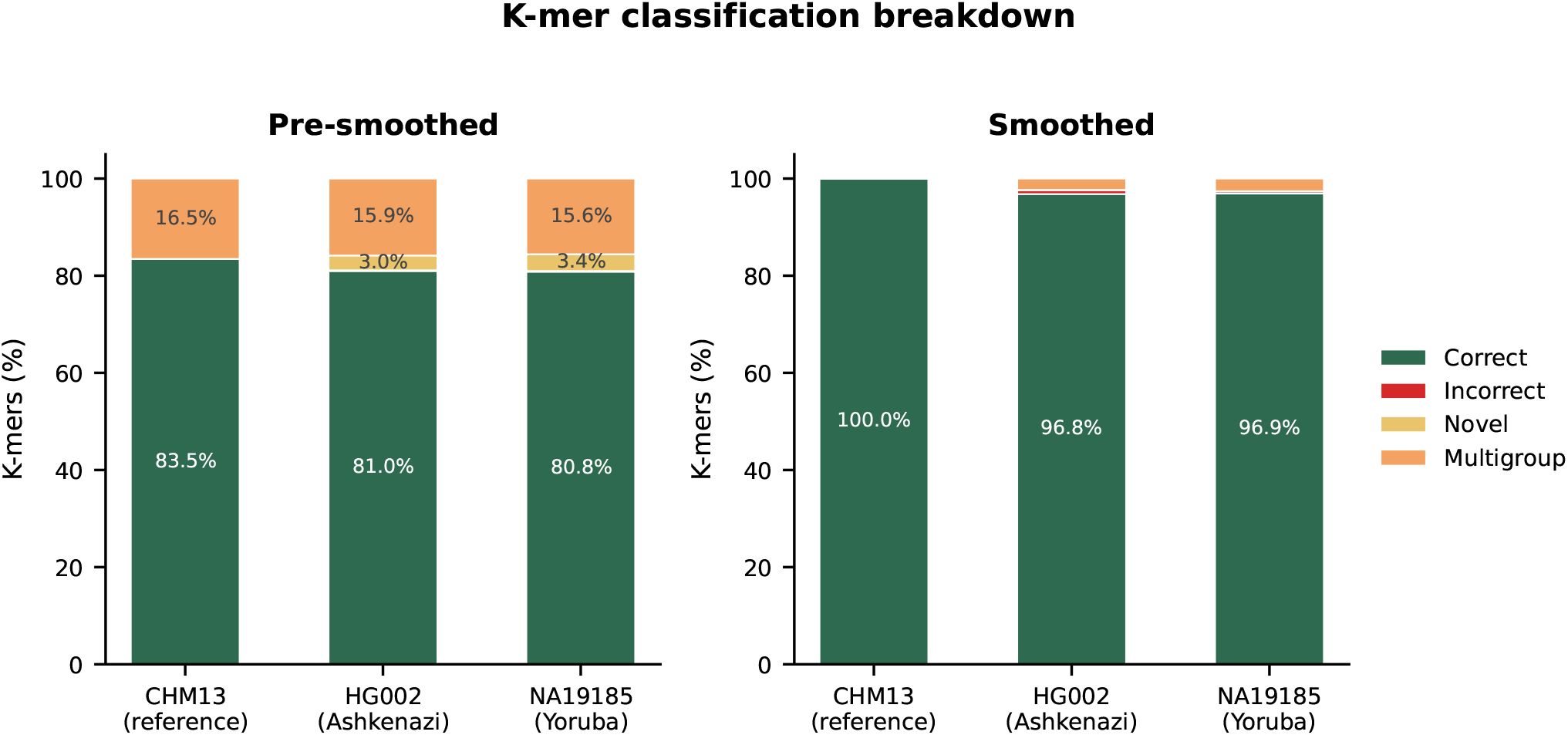
*K*-mer classification before and after smoothing. *K*-mers are categorized as correct (chromosome-specific, matching truth), incorrect (chromosome-specific, not matching truth), novel (absent from the index), or multigroup (assigned to a non-specific hierarchy node). Smoothing resolves the majority of non-specific *k*-mers into correct chromosome-specific assignments.

To investigate misclassifications, we analyzed NA19185. We classified all misclassified *k*-mers by whether the true and predicted chromosomes were both acrocentric (chr13, chr14, chr15, chr21, chr22), a sex chromosome swap (chrX/chrY), or neither. In the smoothed output, 59.9% of all misclassified *k*-mers (16.1M of 26.9M) represent inter-acrocentric swaps, and an additional 5.0% (1.3M) represent sex chromosome swaps attributable to pseudoautosomal regions and other X/Y homologous sequence [24]. These patterns are consistent with well-characterized biological phenomena: the short arms of acrocentric chromosomes undergo frequent non-allelic homologous recombination [15], producing shared sequence blocks that are correctly identified by the *k*-mer index but assigned to a different acrocentric chromosome than the one on which they reside in the query assembly (Figure 4).

**Fig. 4.**
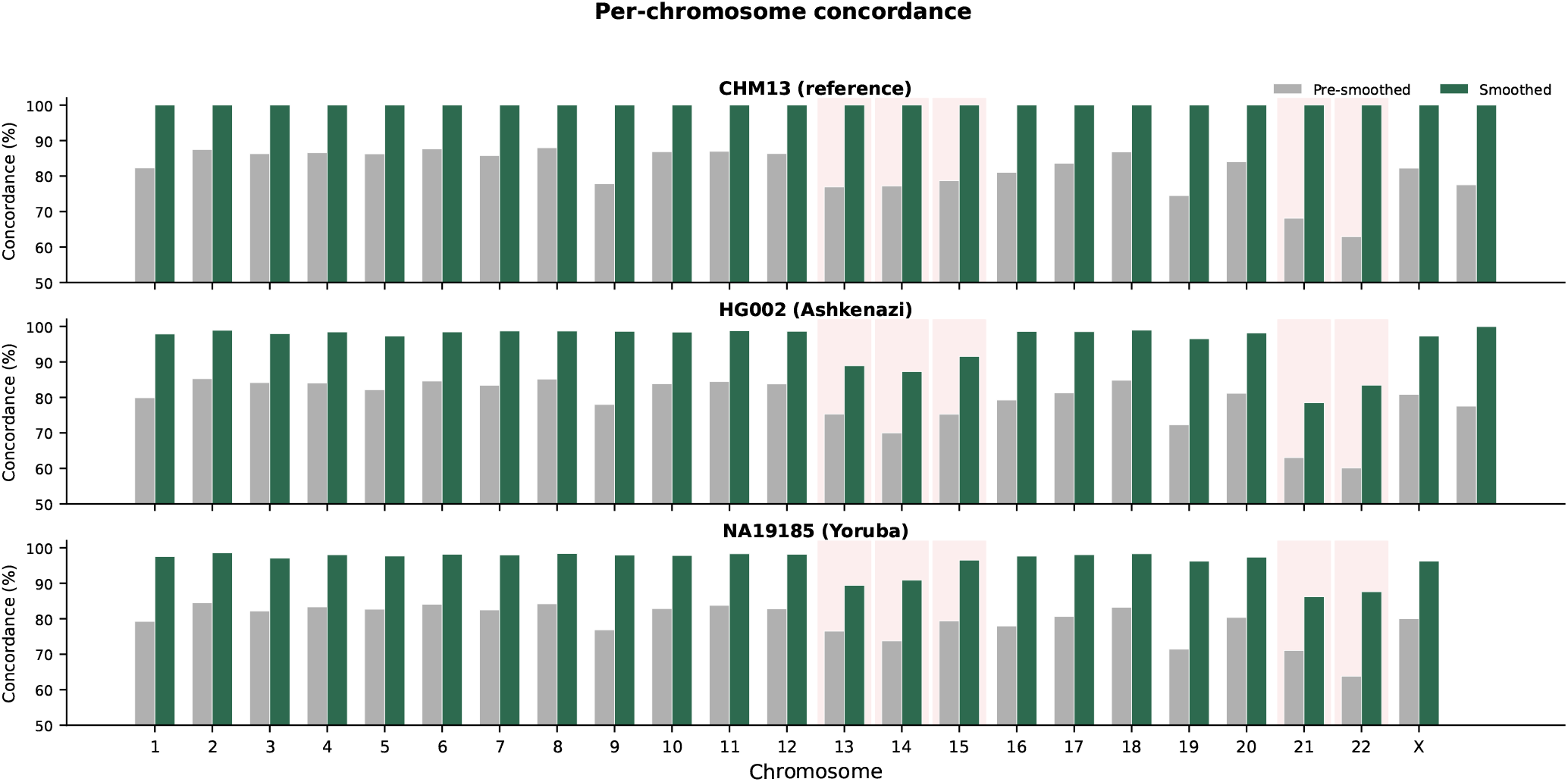
Per-chromosome concordance for pre-smoothed and smoothed annotations. Acrocentric chromosomes (13, 14, 15, 21, 22, shaded) show lower concordance due to inter-chromosomal recombination of their short arms.

The most frequent swap pairs (chr13/chr21, chr14/chr22, chr13/chr15) reflect known patterns of acrocentric short arm sharing. The same pattern explains the apparent accuracy difference between HG002 (99.2%) and NA19185 (99.5%). HG002’s fully assembled 46 T2T scaffolds include more complete acrocentric p-arm sequence, resulting in 41.1M acrocentric swap *k*-mers compared to 16.1M for NA19185 with its 35 T2T scaffolds. After excluding biologically expected mismatches from acrocentric p-arm recombination and sex chromosome homology, the adjusted accuracy is nearly identical: 99.87% for HG002 and 99.84% for NA19185, with adjusted misclassification rates of 0.13% and 0.16%, respectively (Supplementary Figure S3).

The remaining errors (9.4M *k*-mers in NA19185) arise from known structural features: subtelomeric segmental duplications shared between non-homologous chromosomes [23], interior segmental duplications (notably between chromosomes 6 and 7), and pericentromeric repeats. In all cases, these reflect genuine sequence homology rather than algorithmic failures. The smoothing algorithm does not introduce novel error patterns; rather, it amplifies existing biological signals by propagating chromosome labels through non-specific gaps. The smoothing algorithm also handles assembly gaps conservatively: multi-megabase N-gaps in HG002’s acrocentric rDNA arrays are assigned non-specific labels rather than propagating potentially incorrect chromosome-specific labels through the gaps.

### 4.2 Performance comparison

As a performance reference point, we use Kraken2 [35] (referred to as just “Kraken” from here on). Kraken is a highly popular and efficient taxonomic assignment tool for metagenomics, but it can be easily repurposed for any hierarchical *k*-mer assignment task by providing a custom label hierarchy for the index construction procedure. Kraken outputs the most specific labels for each of the input *k*-mers in a run-length coded text format, so the outputs are directly comparable to that of HKS.

The main differences are that HKS supports queries for any *k*-mer length *k* ≤ *s* with the same index, and that Kraken is only approximative in at least three ways:

1. Instead of annotating the *k*-mers, Kraken saves space by annotating *minimizers* of *k*-mers with length *m < k*, leading to potentially false matches if two *k*-mers have the same minimizer.
2. By default, Kraken allows a small number of mismatches in minimizer lookup, increasing sensitivity but potentially reducing accuracy.
3. Kraken uses truncated 32-bit hash keys for indexing minimizers, which introduces the possibility of unresolvable hash collisions. This is a design choice to save space at the expense of rare *k*-mer misclassification errors.

Information loss from (1) can be avoided by setting *m* = *k*, to the detriment of index size and running time. We experiment with various values of *m*. Losses from approximation (2) can be avoided by setting the number of wildcard matches to zero (`--minimizer spaces 0`); we do this in all our experiments to better match the output of HKS. Approximation (3) is inherent in the design of the data structure, there does not seem to be an easy way around it using the available command line flags, so some differences remain in the output. However, despite these differences, the outputs of our tool and Kraken match fairly well (see Figure 5). The strongest signal is found on the diagonal, reflecting contigs in the HG002 assembly (rows) having the highest *k*-mer matches for their respective chromosome in CHM13 (columns). Both tools reveal strong off-diagonal signal among the acrocentric chromosomes (chr13, chr14, chr15, chr21, chr22), consistent with inter-acrocentric short-arm recombination events that occurred in HG002 relative to CHM13 [15].

**Fig. 5.**
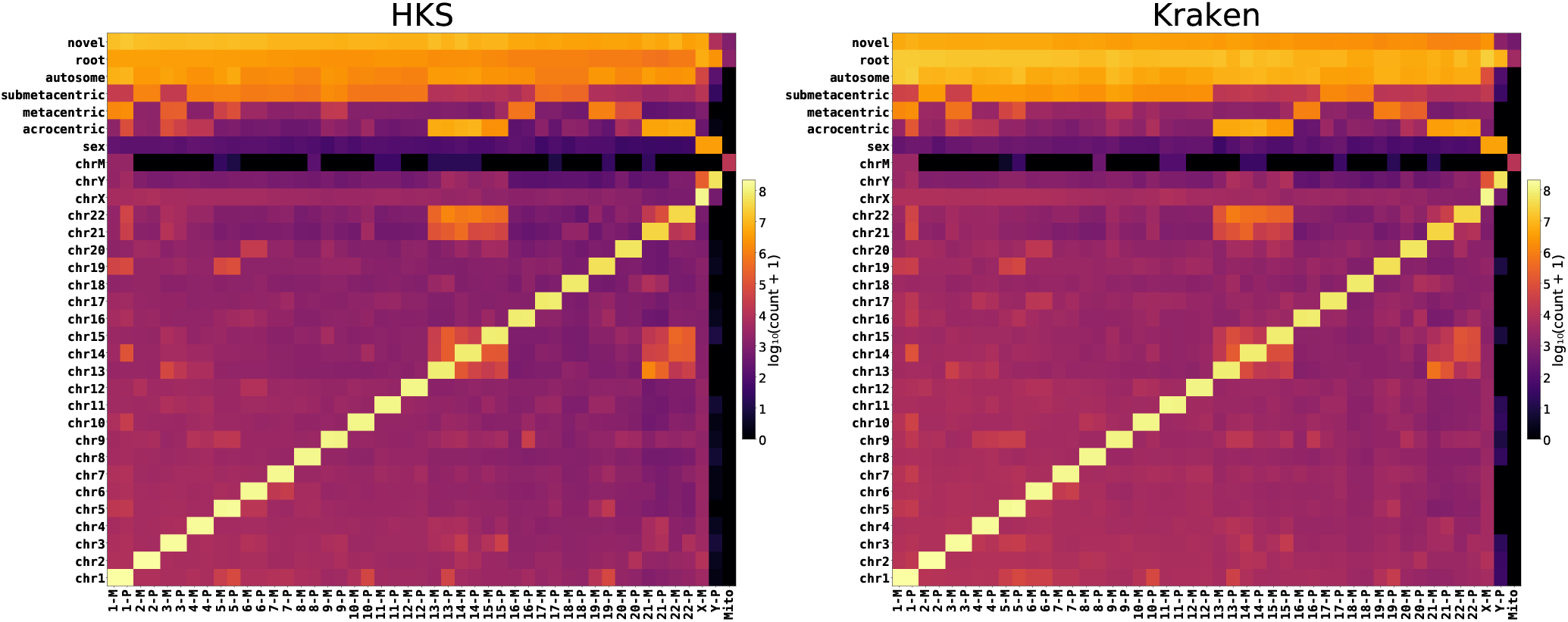
Comparison of HKS (*k* = 63) and Kraken (*k* = 63, *m* = 31, exact minimizer matching) pre-smoothed outputs. The rows are labels in the hierarchy for reference genome CHM13. The columns are chromosomes in the diploid genome HG002 (P for paternal, M for maternal). The color of a cell indicates the number of *k*-mer matching from the input row to the target column, with logarithmic brightness. See Supplementary Figure S5 for the difference of the two heatmaps.

Beyond Kraken, KATKA [14] is a theoretical Lempel-Ziv-based index with *k* given at query time, but no implementation is available. A recent adaptation to maximal exact matches [12] departs significantly from the original

#### Experimental setup

Experiments were conducted on a cluster node running AlmaLinux 8.10 with 503 GiB of RAM and a 64-core AMD EPYC 7302 processor at 3 GHz (32 KiB L1, 512 KiB L2, and 16 MiB L3 cache). Storage was provided by a distributed Lustre file system.

The reference genome is the T2T-CHM13v2.0 haploid human assembly [27], with chromosomes used as categories and the hierarchy shown in Figure 1. The query genome is the diploid human assembly HG002 [16].

#### Performance

We build the HKS index in two stages: in the first stage, we build the SBWT index of the reference data, with *s* = 63, and in the second stage, we add the labels for the *s*-mers in the SBWT. The SBWT construction on CHM13 took 5.5 minutes and 146 GiB of RAM. A disk-based alternative requires only 16 GiB of RAM at the cost of longer construction time (37 minutes) and 93 GiB of temporary disk space. The label annotation phase took 3.2 minutes and used 15.6 GiB of RAM, producing a final index of size 10.4 GiB.

We ran Kraken with all combinations of *k*-mer lengths *k* ∈ {15, 31, 47, 63} and minimizer lengths *m* ∈ {15, 22, 31} with *m* ≤ *k*. Table 3 shows the construction and query throughput compared to that of HKS, using 32 parallel threads. All the HKS queries were run on the single index with *s* = 63, setting the query *k*-mer length equal to the *k*-mer length of the Kraken index. Timing starts when the index is loaded into memory and ready for use. We also include the *s*-mer preprocessing in the index loading time as it is a linear-time scan that can be performed concurrently as loading the index.

**Table 3.** Comparison of Kraken (with minimizer length *m* ≤ *k*, written as Kraken-m) and hks at *t* = 32 threads and varying *k*-mer length *k*. All hks results use the same index built for maximum query length *s* = 63. Columns: build throughput (*B*_*t*_, megabases / second), peak memory during build (*B*_*m*_, GiB), index size (*I*, GiB), and query throughput (*Q*_*t*_, megabases / second). hks construction proceeds in two phases: first, an SBWT index is built in memory, and second, the hks index is constructed from the SBWT. The *B*_*t*_ and *B*_*m*_ values reported for hks are for the whole two-stage pipeline. The second phase took 189 seconds and used 15.6 GiB of RAM. A slower disk-based option exists for SBWT construction, taking 37 minutes, using 15.9 GiB of RAM and 92.9 GiB of disk.

| | $k = 15$ | | | | $k = 31$ | | | | $k = 47$ | | | | $k = 63$ | | | |
| --- | --- | --- | --- | --- | --- | --- | --- | --- | --- | --- | --- | --- | --- | --- | --- | --- |
| | $B_t$ | $B_m$ | $I$ | $Q_t$ | $B_t$ | $B_m$ | $I$ | $Q_t$ | $B_t$ | $B_m$ | $I$ | $Q_t$ | $B_t$ | $B_m$ | $I$ | $Q_t$ |
| Kraken-m15 | 2.5 | 3.2 | 1.8 | 66.6 | 12.8 | 3.2 | 0.3 | 133.9 | 17.9 | 3.2 | 0.2 | 138.8 | 21.1 | 3.1 | 0.1 | 140.8 |
| Kraken-m22 | NA | NA | NA | NA | 6.6 | 3.2 | 2.2 | 130.1 | 13.5 | 3.2 | 0.9 | 137.4 | 18.0 | 3.2 | 0.6 | 138.7 |
| Kraken-m31 | NA | NA | NA | NA | 1.3 | 14.1 | 13.4 | 69.7 | 10.1 | 3.2 | 1.5 | 136.1 | 14.9 | 3.2 | 0.8 | 138.5 |
| HKS | 9.4 | 146.0 | 10.4 | 22.0 | 9.4 | 146.0 | 10.4 | 83.7 | 9.4 | 146.0 | 10.4 | 86.3 | 9.4 | 146.0 | 10.4 | 85.3 |

#### Queries

We find that HKS provides comparable query throughput compared to Kraken, despite Kraken using a specialized index for each length *k*, and using lossy methods to improve query time. When the minimizer length *m* is set equal to *k*, such as *k* = *m* = 31, the throughput of Kraken drops by more than half. In contrast, HKS maintains consistent query performance for all query lengths except for *k* = 15, where our single-threaded output formatting becomes a bottleneck. Regarding parallelism, for *k >* 15, we observe close to ideal parallel query speedup when using up to 16 threads, but beyond that, the performance starts to plateau (Supplementary Figure S4). The CPU utilization remains high, however. Since our lookup algorithm is likely memory-access bound due to its unpredictable memory access pattern, this suggests that our implementation might be close to saturating the maximum random-access throughput of the memory hardware.

#### Index size

The HKS index for *s* = 63 takes 10.4 GiB on disk. The index size of Kraken is highly dependent on the value of *m*. The larger the gap between *k* and *m*, the smaller the index. At the most extreme data point of *k* = 63, *m* = 15, the Kraken index is over a hundred times smaller than HKS. However, this comes at a significant loss of information, and at more typical parameters such as *k* = 31, *m* = 22, the difference is only five-fold, and at the other extreme of *k* = 31, *m* = 31, the HKS index is slightly smaller (10.4 GiB vs 13.4 GiB). Parameter values with *m >* 31 could not be experimented with since Kraken supports only *m* up to 31.

We emphasize that despite HKS having a larger index, it is significantly more flexible and accurate as it essentially contains a lossless version of *every* Kraken index for *k* from 1 to 63. This allows the user to make an informed decision for which query *k*-mer length to use in their application. To aid with this choice, HKS provides a subcommand to count the number of *k*-mers associated with each node for each 1 ≤ *k* ≤ *s*. Figure 6 illustrates the resulting output for our experimental dataset. We see that the number of nodes assigned to the root peaks at around *k* = 15, and by *k* = 20, most labels have been pushed down to the leaves and the assignment ratios start to stabilize.

**Fig. 6.**
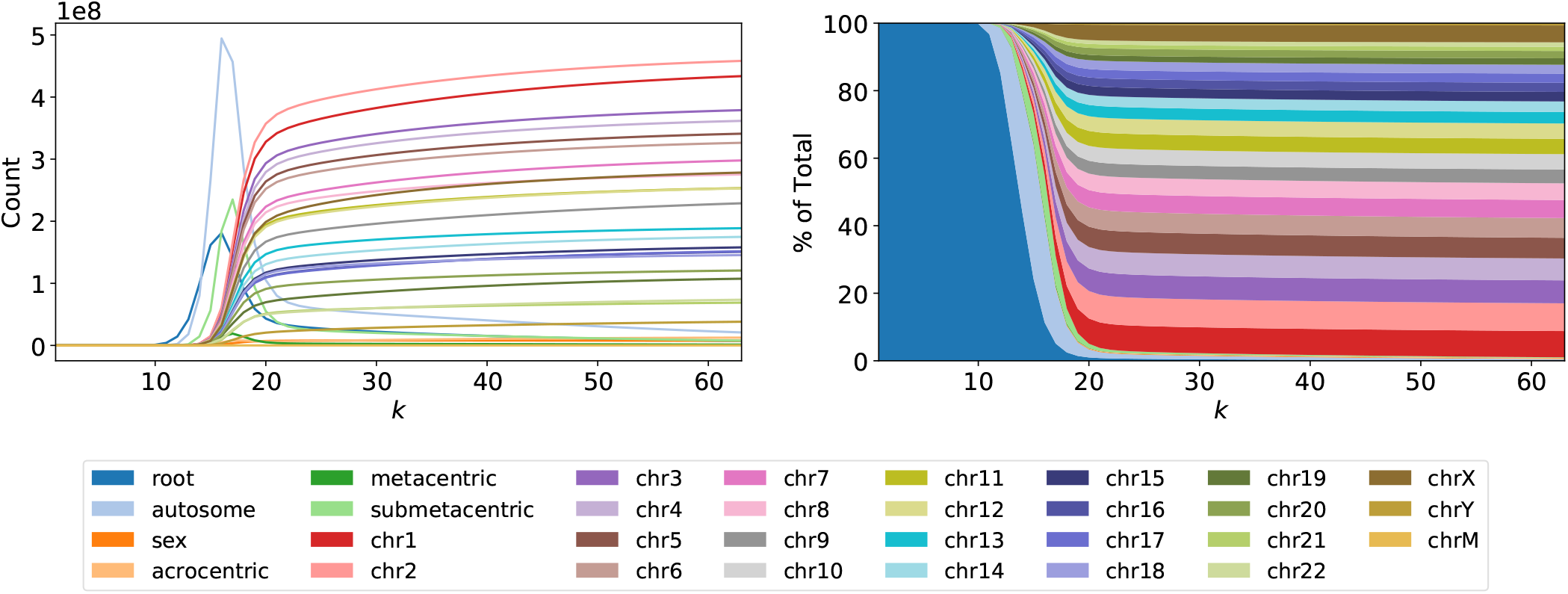
Left: count of *k*-mers for 1 ≤ *k* ≤ 63 assigned to each node label in the hierarchy. Right: the fraction of *k*-mers assigned to each node label in the hierarchy. Kraken approach. Generic colored de Bruijn graph tools such as Themisto [6], Fulgor [13], and Metagraph [20] could in principle be used for hierarchical *k*-mer annotation by post-processing color sets or by precomputing hierarchy labels as colors. However, these tools are not specifically optimized for hierarchical labeling and offer many alternative indexing schemes, so we exclude them from our benchmarks.

## 5 Conclusion

We have presented HKS, a data structure for exact hierarchical *k*-mer annotation that supports variable-length queries from a single index. By augmenting the Spectal Burrows Wheeler Transform with entry points into a labeled hierarchy, HKS achieves query throughput comparable to Kraken2 while providing lossless annotation across all *k*-mer lengths from 1 to *s*. The feature assignment framework formalizes how indexed *k*-mers can be partitioned into disjoint sets according to a user-defined category hierarchy, and the hierarchy-aware smoothing algorithm recovers chromosome-specific assignments for ∼ 97% of query *k*-mers, with residual errors attributable to well-characterized biological phenomena rather than algorithmic failures.

Several directions for future work remain. For example, the information stored in the HKS index could be used to automatically select the best value of *k*, potentially improving accuracy and removing one manually tuned parameter. The current validation uses a single haploid reference genome (T2T-CHM13v2.0); extending the approach to pangenome references could improve sensitivity for diverse populations by reducing the fraction of novel *k*-mers. While we have demonstrated the approach on chromosome and repeat feature sets, the framework is general and could be applied to other hierarchical annotation tasks such as taxonomic profiling or transcript-level quantification. Finally, the smoothing algorithm currently operates on each feature set independently; jointly leveraging information across feature sets — for example, using repeat annotations to inform chromosome assignment at repeat-dense boundaries — could further improve specificity.

## Acknowledgments

This study was funded by the WILL Chair BOSSA 2025-2029 of University of Lille.

