## Supplementary information for "Hierarchical genomic feature annotation with variable-length queries"

### Supplementary Materials

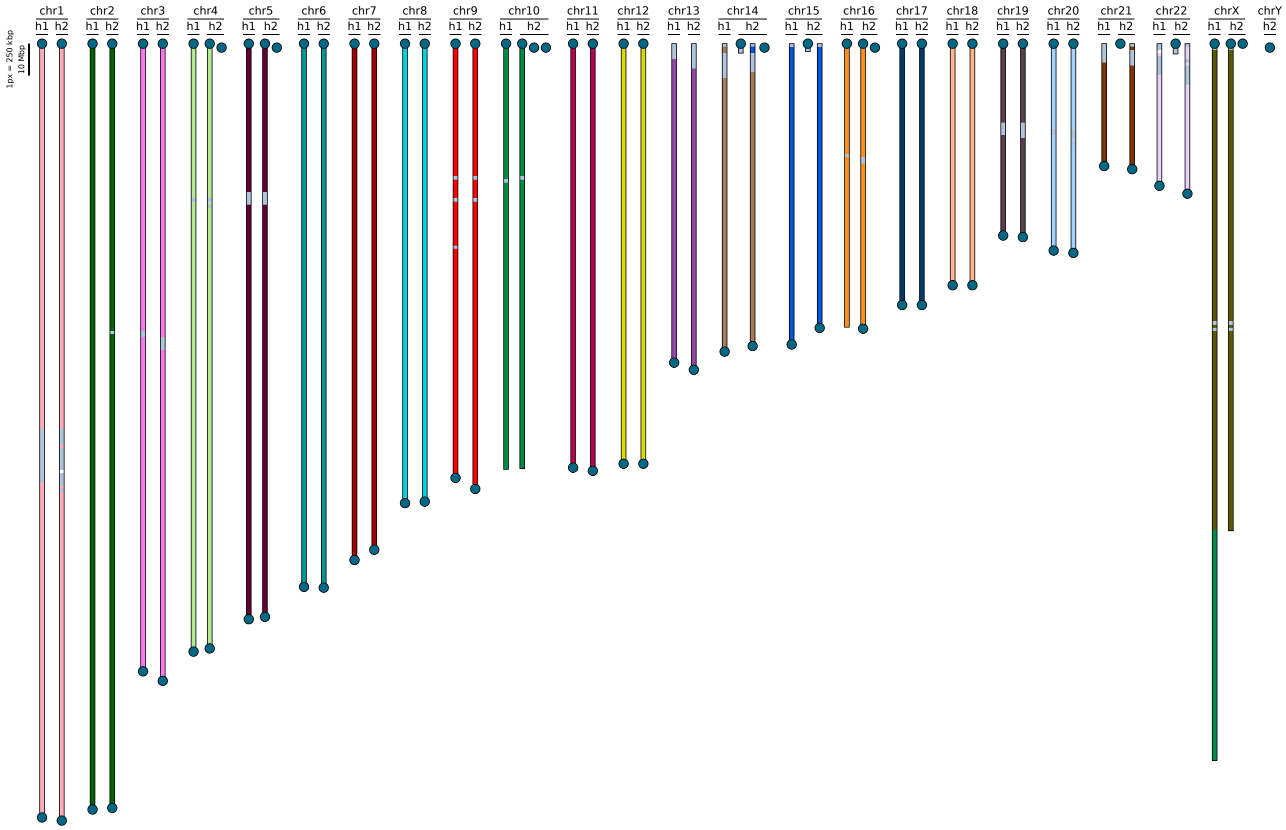


Fig. S1: Positional annotation of the diploid RPE1 genome assembly produced by HKS using chromosomes as categories, derived from T2T-CHM13v2.0. Each $k$-mer in each contig is assigned a chromosomal label (indicated by bar color), and consecutive $k$-mers sharing the same label are merged into intervals. Intervals are drawn proportional to genomic length (scale bar, top left). Contigs are assigned to chromosomes (column headers) based on majority $k$-mer assignment, with haplotypes denoted by h1 and h2. A blue circle indicates that a contig terminates in a telomere and contigs marked by telomeres on both ends most likely represent telomere-to-telomere chromosomes. Segments of discordant color near contig termini (on chromosomes 13-15,21,22,X) or centromeric regions (most other chromosomes) reflect pseudohomologous and satellite-derived $k$-mers not unique to any single chromosome. The segmentation reveals a translocation between chrX and chr10, visible as a chr10-colored (green) interval within the chrX h1 contig.

| Contig | Haplotype | Accession |
|---|---|---|
| NA19185#1#JBHDXR010000014.1 | 1 | JBHDXR010000014.1 |
| NA19185#1#JBHDXR010000050.1 | 1 | JBHDXR010000050.1 |
| NA19185#1#JBHDXR010000052.1 | 1 | JBHDXR010000052.1 |
| NA19185#2#JBHDXS010000037.1 | 2 | JBHDXS010000037.1 |
| NA19185#2#JBHDXS010000038.1 | 2 | JBHDXS010000038.1 |
| NA19185#2#JBHDXS010000039.1 | 2 | JBHDXS010000039.1 |
| NA19185#2#JBHDXS010000040.1 | 2 | JBHDXS010000040.1 |

Table S1: Acrocentric p-arm contigs in the NA19185 assembly excluded from the chromosome assignment benchmark. These contigs could not be confidently assigned to a specific acrocentric chromosome of origin due to extensive sequence sharing among acrocentric short arms.

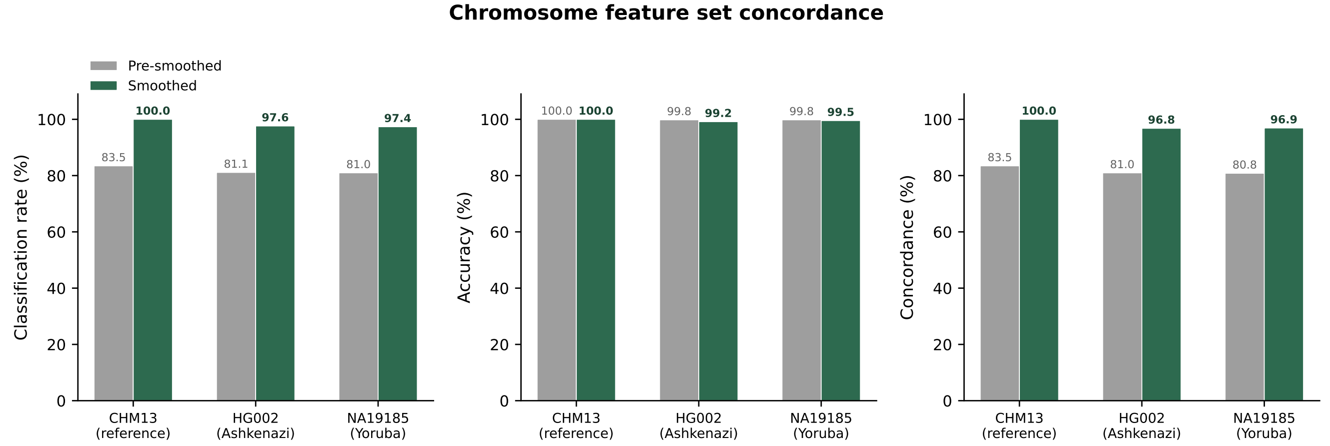


Fig. S2: Chromosome feature set concordance across three genome assemblies. Classification rate, accuracy, and overall concordance are shown for pre-smoothed and smoothed annotations.

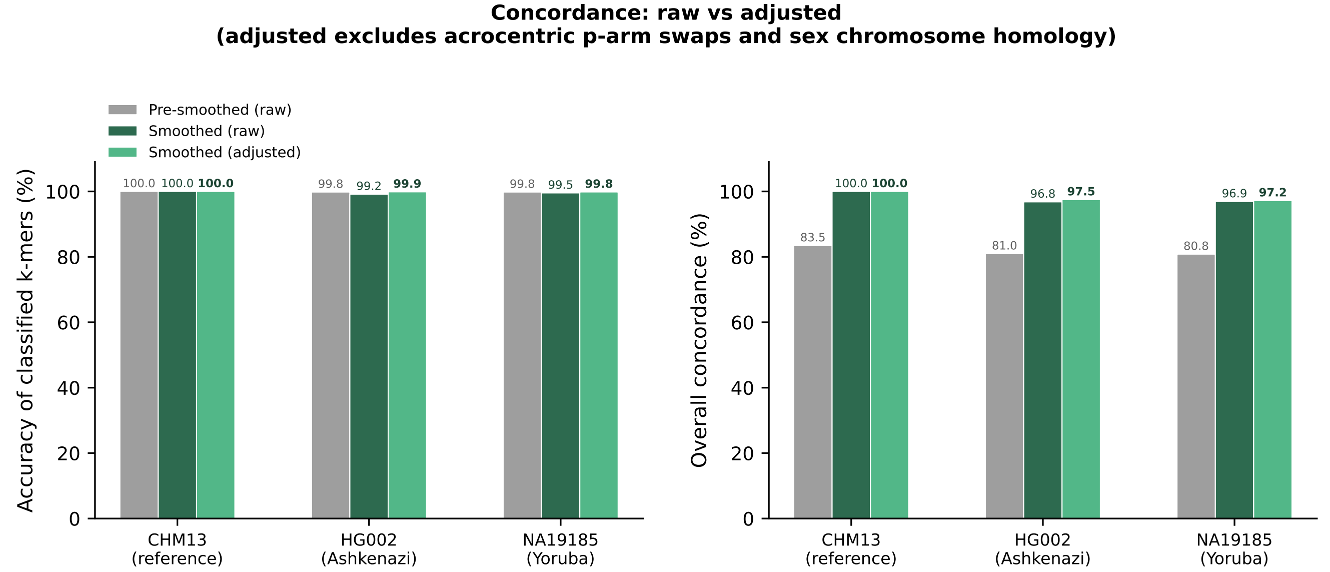


Fig. S3: Raw versus adjusted accuracy and concordance. Adjusted metrics exclude inter-acrocentric chromosome swaps and sex chromosome homology. The apparent accuracy difference between HG002 and NA19185 disappears after adjustment, indicating it is driven by differences in acrocentric p-arm assembly completeness.

**Algorithm S1** Hierarchy-aware smoothing

**Require:** Ordered intervals $I = [i_1, i_2, \ldots, i_n]$ with features $f(i_j)$; category hierarchy $H$; gap threshold $g$
**Ensure:** Smoothed feature assignments

```
repeat
    changed ← false
    for each anchor i_j in I do
    // Ancestor phase
        t ← j;  f_last ← f(i_j);  disallowed ← ∅
        for m = j+1 to n do
            if gap(i_m, i_t) > g then
                break
            end if
            if f(i_m) is ancestor of f_last in H and f(i_m) ∉ disallowed then
                t ← m;  f_last ← f(i_m)
            else if f(i_m) is unrelated to f_last then
                disallowed ← disallowed ∪ ANCESTORS(f(i_m))
            else
                break
            end if
        end for
    // Descendant phase
        p ← f(i_t);  r ← t
        for m = t+1 to n do
            if gap(i_m, i_r) > g then
                break
            end if
            if f(i_m) is descendant of f_last in H then
                r ← m;  f_last ← f(i_m)
            else if f(i_m) is unrelated to f_last and p ∈ ANCESTORS(f(i_m)) then
                break
            else if f(i_m) is ancestor of f_last then
                break
            end if
        end for
    // Reassignment
        if r > j then
            ℓ ← LCA(f(i_j), f(i_r))
            for each a, j < a < r do
                if ℓ is ancestor of f(i_a) in H then
                    f(i_a) ← ℓ;  changed ← true
                end if
            end for
        end if
    end for
until ¬changed
```

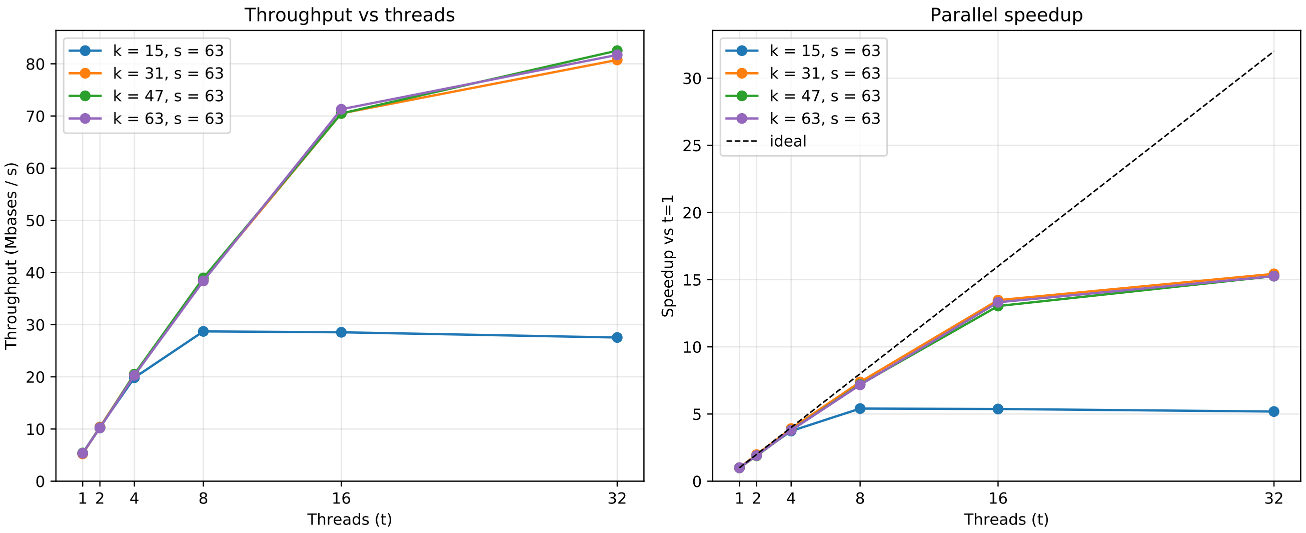


Fig. S4: Performance of $k$-mer queries on a 63-mer index of T2T-CHM13v2.0 as a function of the number of threads. The query consists of both copies of chromosome 1 in the diploid human reference genome HG002. The query for $k = 15$ has a significantly larger output file, so output formatting becomes a bottleneck, and the query throughput saturates earlier.

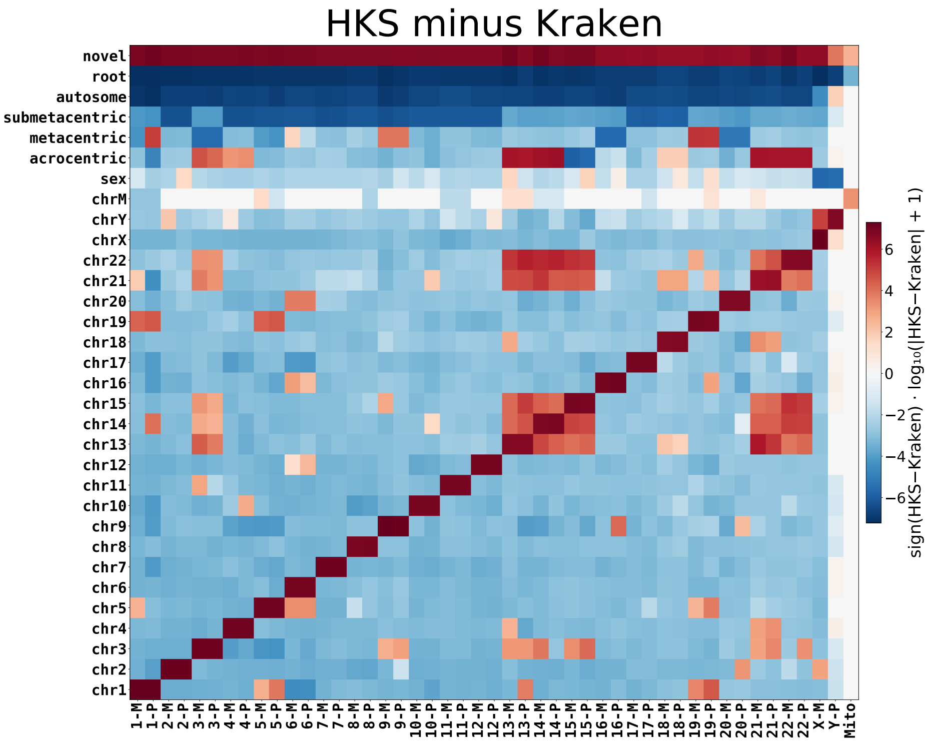


Fig. S5: Difference between HKS ($k = 63$) and Kraken ($k = 63, m = 31$) outputs in Figure 5.

| **Repeat category** | **Priority** |
|---|---|
| D20S16 | 1 (highest) |
| rRNA | 2 |
| scRNA | 3 |
| snRNA | 4 |
| tRNA | 5 |
| RC | 6 |
| Retroposon | 7 |
| DNA | 8 |
| LTR | 9 |
| SINE | 10 |
| LINE | 11 |
| Unknown | 12 (lowest) |

Table S2: Priority order for resolving overlapping RepeatMasker annotations. When a genomic position is annotated with multiple repeat categories, the category with the highest priority (lowest number) is retained. Positions with no RepeatMasker annotation are assigned to the `nonrepeat` category.
